## Supplementary Information for "From Hi-C Contact Map to Three-dimensional Organization of Interphase Human Chromosomes"

Guang Shi[1] and D. Thirumalai[1*]

[1] *Department of Chemistry,*

*University of Texas at Austin, Texas, 78712*

(Dated: May 19, 2020)

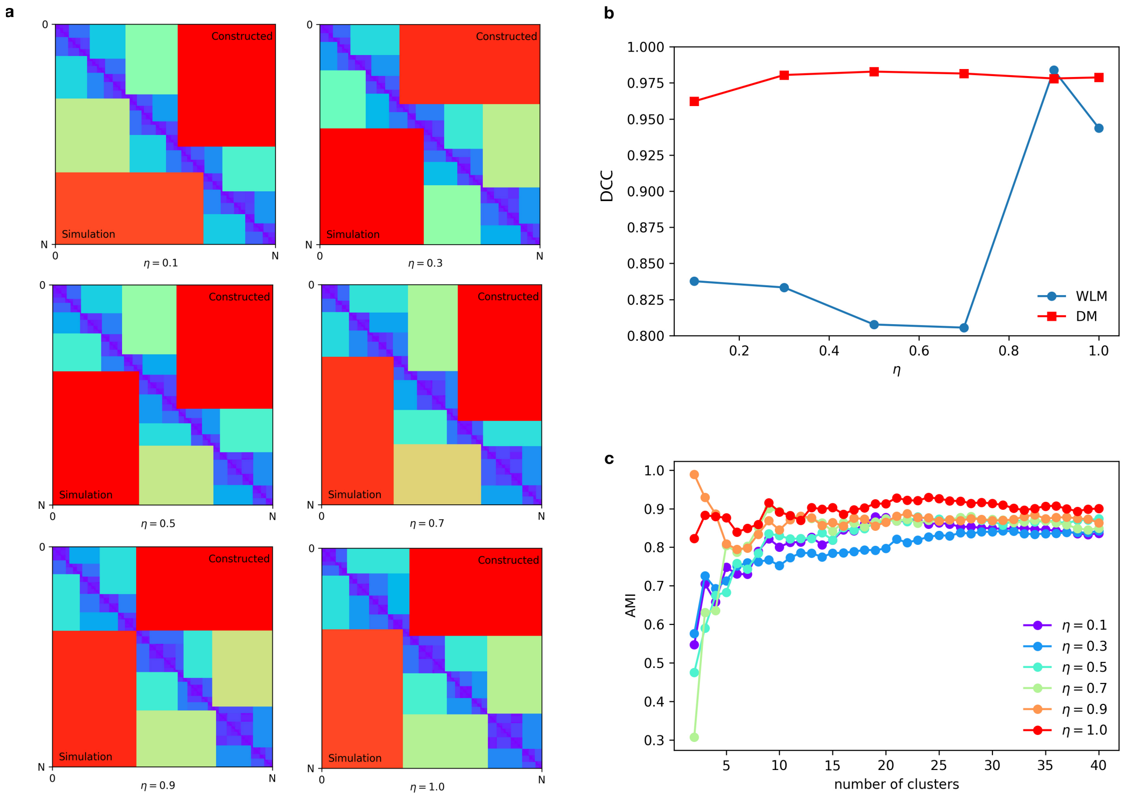


FIG. S1: **(a)** Comparison between simulated and constructed Ward Linkage Matrices (WLMs) for $\eta = (0.1, 0.3, 0.5, 0.7, 0.9, 1.0)$. WLMs are constructed from the mean distance matrices (DMs) for GRMC. For each subgraph, the simulated WLM and the constructed WLM are plotted in lower and upper triangle, respectively. **(b)** Distance Correlation coefficients (DCC) as a function of $\eta$. Blue and red dots are for DCC computed using WLMs and DMs, respectively. Interestingly, the DCC is high ($> 0.8$) even for small $\eta$. **(c)** Adjusted Mutual Information (AMI) score used to quantify the hierarchical clustering as a function of number of clusters.

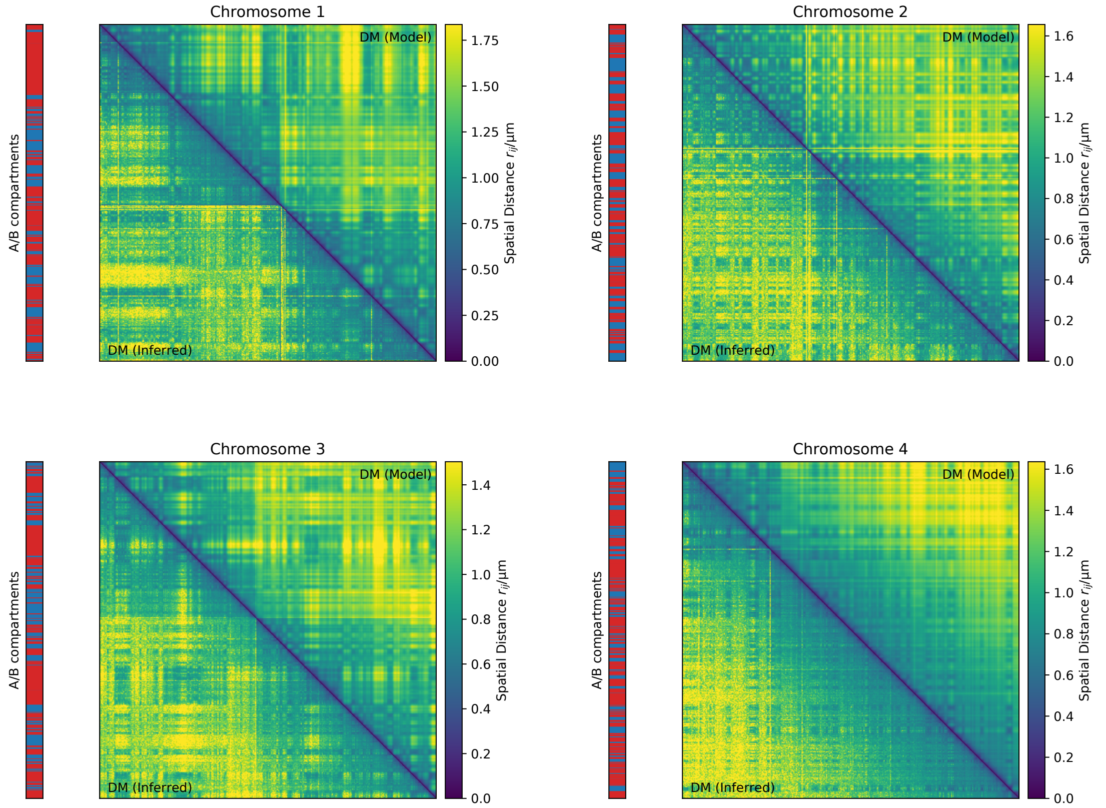


FIG. S2: Comparison between the DM inferred from Hi-C data (lower triangle) and the DM calculated from an ensemble of 3D structures for Chromosomes 1-4 using the HIPPS method, $\langle \mathbf{P} \rangle \rightarrow \langle \bar{\mathbf{R}} \rangle \rightarrow$3D structures (upper triangle)

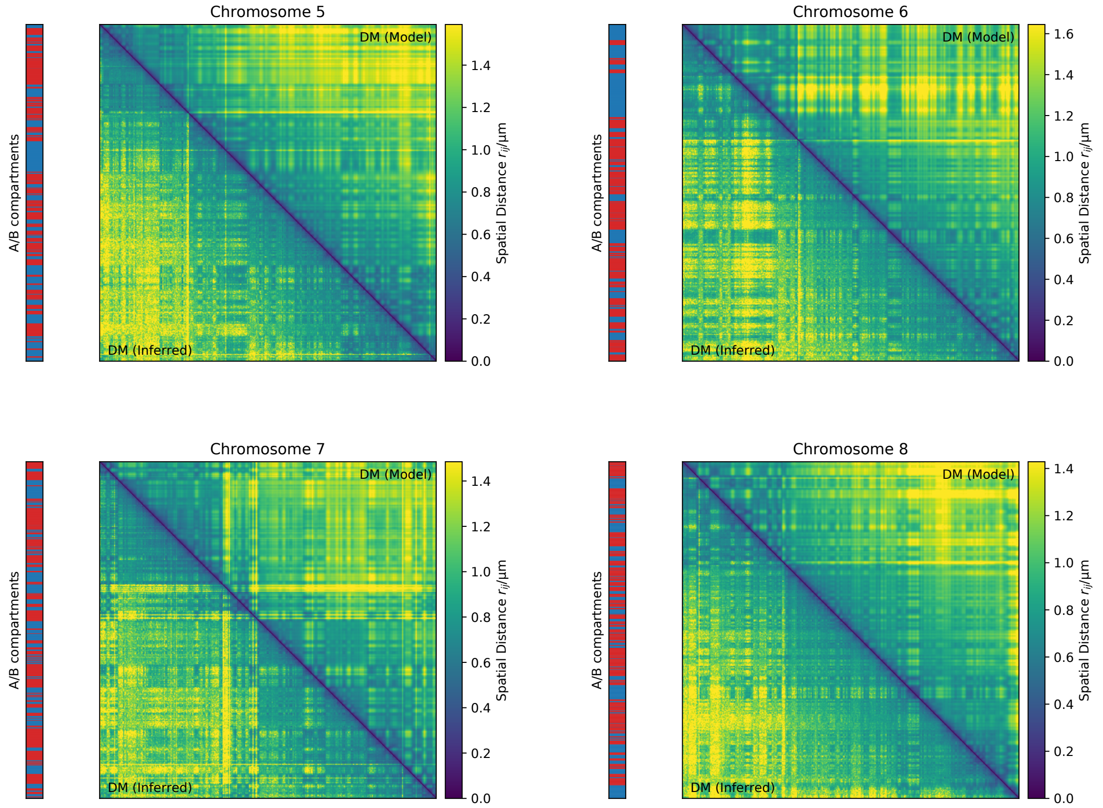


FIG. S3: Same as Fig. S2 except these are for Chromosomes 5-8.

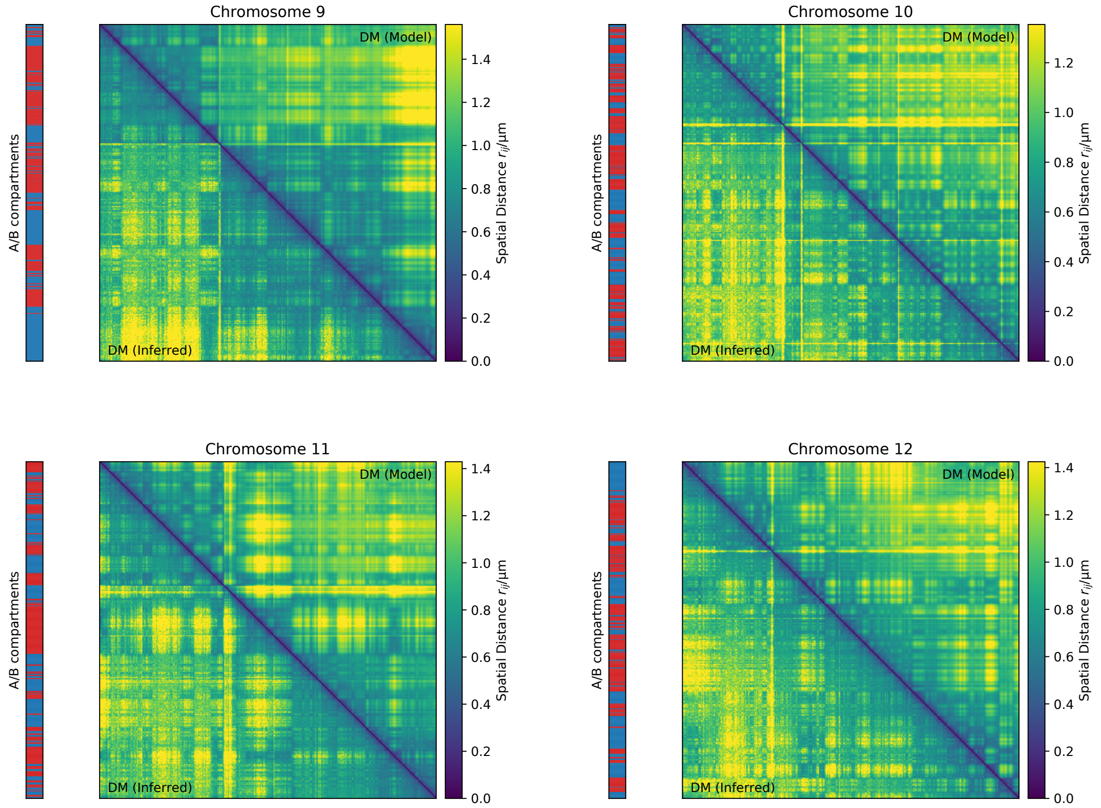


FIG. S4: Same as Fig. S2 except these are for Chromosomes 9-12.

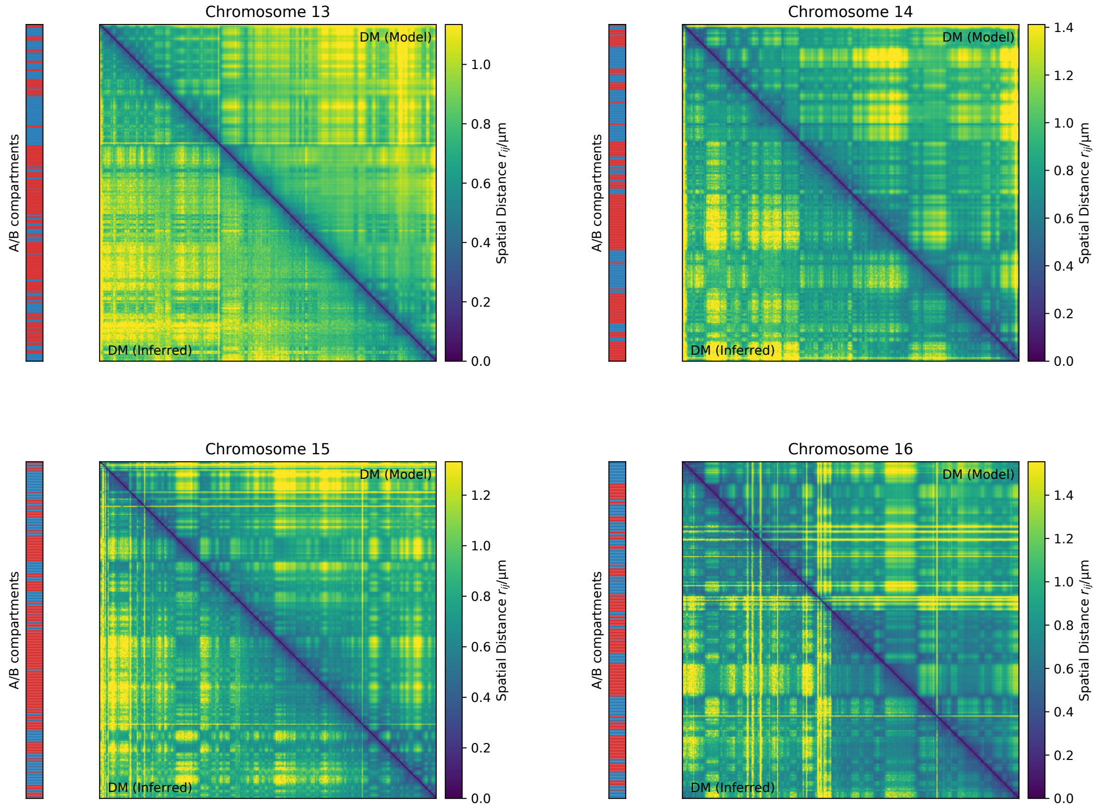


FIG. S5: Same as Fig. S2 except these are for Chromosomes 13-16.

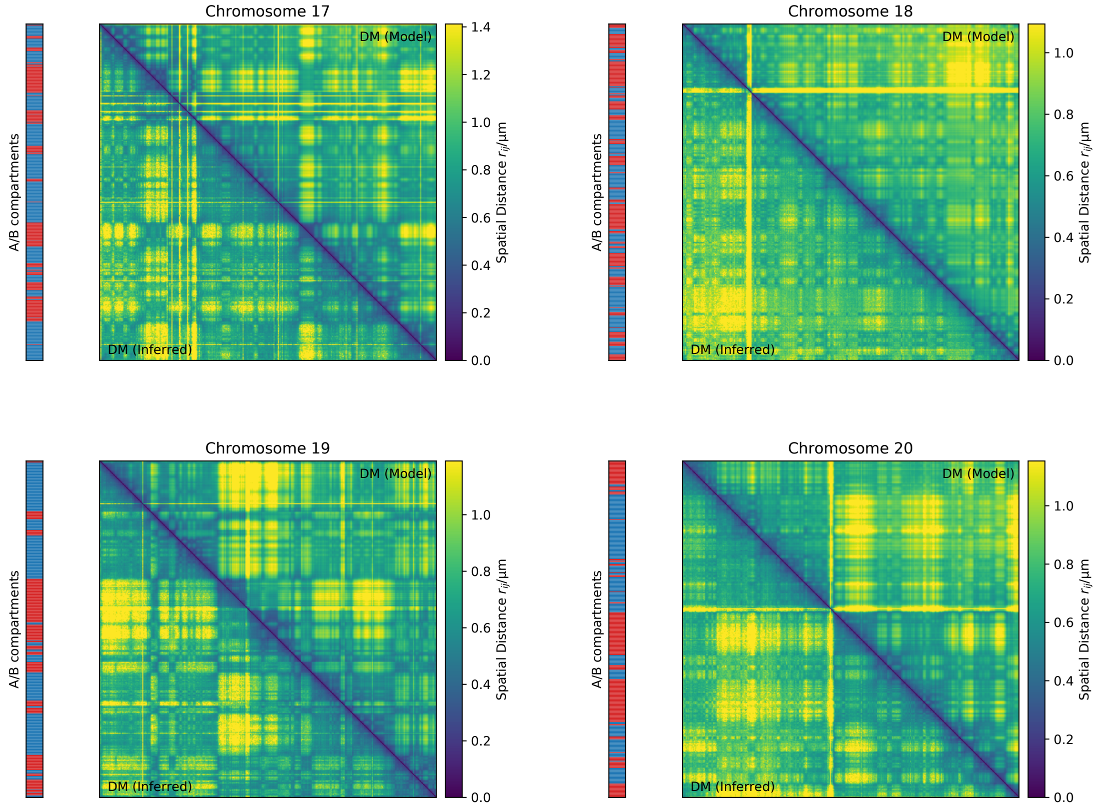


FIG. S6: Same as Fig. S2 except these are for Chromosomes 17-20.

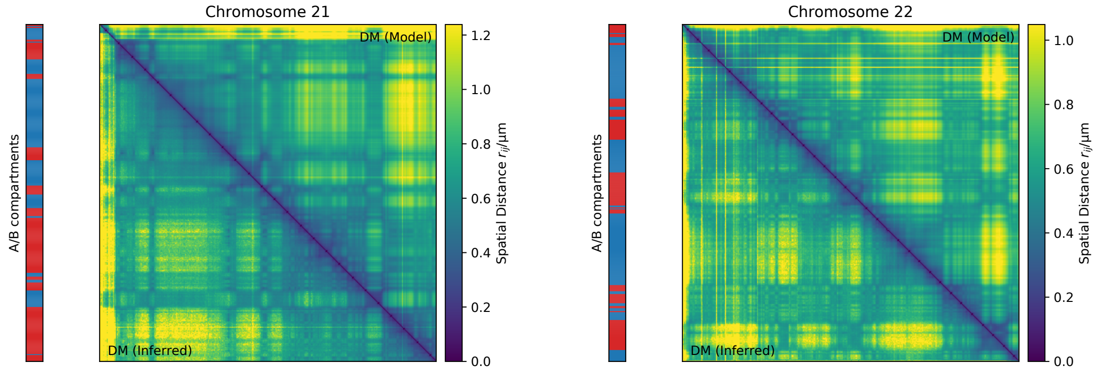


FIG. S7: Same as Fig. S2 except these are for Chromosomes 21 and 22.

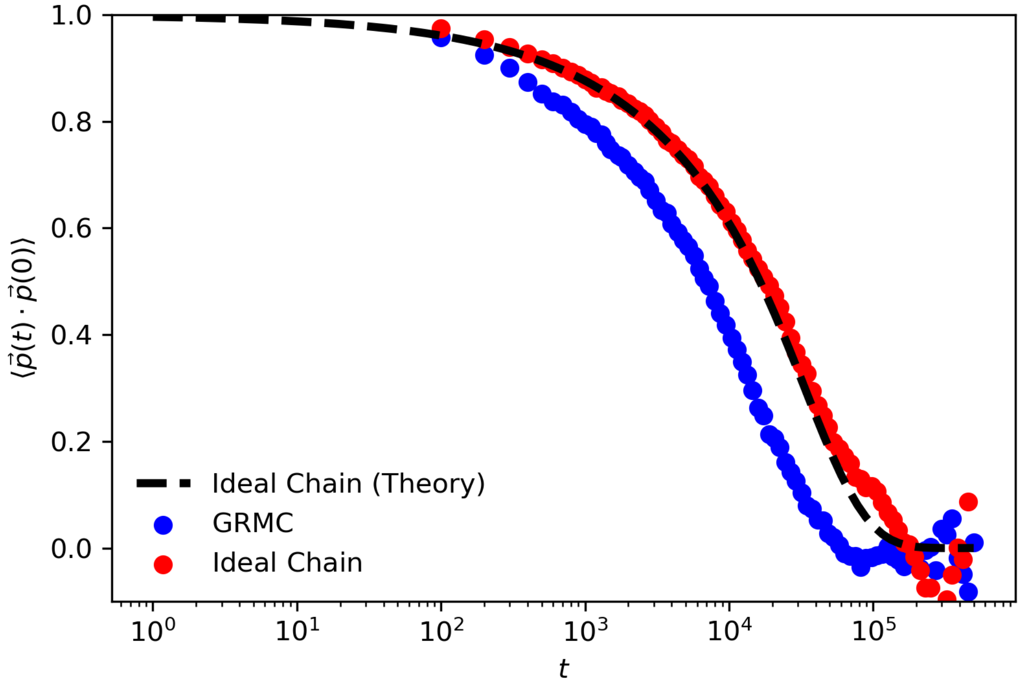


FIG. S8: Autocorrelation of the end-to-end vector. $\vec{p}(t)$ is the end-to-end vector at time $t$. The dashed curve is the theoretical prediction for ideal chain from Rouse model. Our simulation of ideal chain agrees with the theory as expected. The relaxation time $\tau$ (when $\langle \vec{p}(t) \cdot \vec{p}(0) \rangle = 1/e$) for ideal chain and GRMC is about $3 \times 10^6$ timesteps and $10^6$ timesteps, respectively. Thus our simulation produces about 800 independent snapshots for GRMC and 300 independent snapshots for ideal chain.

| Chr | $N_c$ | Chr | $N_c$ | Chr | $N_c$ |
|---|---|---|---|---|---|
| Chr1 | 2225 | Chr9 | 1090 | Chr17 | 777 |
| Chr2 | 2374 | Chr10 | 1294 | Chr18 | 750 |
| Chr3 | 1948 | Chr11 | 1309 | Chr19 | 558 |
| Chr4 | 1875 | Chr12 | 1306 | Chr20 | 599 |
| Chr5 | 1760 | Chr13 | 960 | Chr21 | 344 |
| Chr6 | 1676 | Chr14 | 879 | Chr22 | 338 |
| Chr7 | 1542 | Chr15 | 797 | ChrX | 1509 |
| Chr8 | 1414 | Chr16 | 784 | | |

TABLE S1: Values of $N_c$, the number of loci, for the 23 chromosomes.
